# A Nanovial-Based Platform for Functional Discovery of Antigen-Reactive TCRs from Unconventional T Cells

**DOI:** 10.1101/2025.08.06.668816

**Authors:** Citradewi Soemardy, Yan-Ruide Li, Yichen Zhu, Lili Yang, Dino Di Carlo

**Affiliations:** Department of Bioengineering, University of California Los Angeles, Los Angeles, California 90095, USA; Department of Microbiology, Immunology & Molecular Genetics, University of California, Los Angeles, Los Angeles, CA 90095, USA; Jonsson Comprehensive Cancer Center, University of California Los Angeles, Los Angeles, California 90095, USA; Eli and Edythe Broad Center of Regenerative Medicine and Stem Cell Research, University of California, Los Angeles, Los Angeles, CA 90095, USA; Jonsson Comprehensive Cancer Center, David Geffen School of Medicine, University of California, Los Angeles, Los Angeles, CA 90095, USA; Parker Institute for Cancer Immunotherapy, University of California, Los Angeles, Los Angeles, CA 90095, USA; California NanoSystems Institute (CNSI), University of California Los Angeles, Los Angeles, California 90095, USA

**Author notes:** Co-first authors (Citradewi Soemardy, Yan-Ruide Li, Yichen Zhu).

## Abstract

Unconventional T cells, such as mucosal-associated invariant T (MAIT) cells and invariant natural killer T (iNKT) cells, recognize non-peptide antigens presented by MR1 and CD1d, respectively, and play pivotal roles in immunity, allowing targeting of cells based on metabolic activity. Although components of the T cell receptors (TCRs) for these unconventional T cells are invariant, significant variability in the CDR3 regions still exist, opening questions as to how TCR sequence and function may be linked, and how to maximize the therapeutic potential of engineered unconventional T cells. Here, we develop a nanovial-based functional screening platform that enables high-throughput discovery of TCRs from unconventional T cells based on direct antigen recognition and cytokine secretion. By selectively labeling nanovials with MR1 and CD1d molecules displaying their cognate ligands, we achieve dose-dependent capture and activation of MAIT and iNKT cells from complex human PBMC samples comprising tens of millions of cells. Using oligonucleotide barcodes conjugated to nanovials encoding the antigen-presenting molecules and loading cytokine capture antibodies, we perform secretion-encoded single-cell sequencing to link TCR identity, gene expression, antigen specificity, and functional response. Applying this method, we isolate rare reactive T cells, recover their TCRs, and validate five novel MAIT TCRs. All five TCRs, when re-expressed in primary T cells, confer antigen-specific cytokine secretion and cytotoxicity. The top two TCRs were evaluated using an in vivo solid tumor model, demonstrating specific tumor homing and efficacy. This function-first strategy offers a powerful tool to uncover functional TCRs from unconventional T cells, yielding a 100% hit rate when secretion-based validation is included as part of the initial screen, unlocking new opportunities for cell-based immunotherapy.

## Introduction

Immunotherapies that harness the antigen-specific activity of T cells have revolutionized cancer treatment, offering durable responses across multiple malignancies^1,2^. While current strategies such as checkpoint blockade and engineered T cell therapies have predominantly focused on conventional peptide-MHC–restricted αβ T cells, there is growing recognition of the unique and often untapped potential of unconventional T cells—including mucosal-associated invariant T (MAIT) cells, invariant natural killer T (iNKT) cells, and γδ T cells^3–8^. These cells recognize specific non-peptide antigens presented by non-classical MHC-like molecules such as MR1 and CD1d and possess distinct effector properties that may be particularly suited to targeting tumors in immunosuppressive microenvironments^3–5,8^.The lack of classical HLA restriction also opens the possibility for development of allogeneic, “off-the-shelf” therapeutic strategies without the risk of graft-versus-host disease (GvHD).

Among these, MAIT cells represent a promising but underexplored subset. They express a semi-invariant TCR typically composed of a TRAV1-2/TRAJ33 α-chain and respond to microbially-derived riboflavin metabolites presented by MR1^5,9^. Notably, recent studies have identified MAIT cell activity in tumor contexts, and increased expression of MR1 in some tumors, suggesting broader roles beyond microbial immunity^6,9–11^. MAIT cells are capable of targeting both MR-1 expressing tumor cells through cognate recognition and MR-1-negative tumor cells via natural killer (NK)-like mechanisms^6^. As such, MAIT cells offer a unique set of advantages for cancer immunotherapy, including a universal antigen restriction, innate tissue-homing to mucosal sites, rapid effector functions, and low alloreactive potential^6,12–14^.

Similarly, iNKT cells have also shown promise in the context of cancer immunotherapy. In humans, they express the TRAV10/TRAJ18 (Vα24-Jα18) invariant TCR α-chain paired with a limited set of TCRβ chains^4,15–19^. iNKT cells are known to recognize glycolipid antigens α-galactosylceramide (αGalCer) presented by the monomorphic CD1d molecules^16^. Recent studies have demonstrated that iNKT cells can directly kill CD1d-expressing tumor cells and remodel the tumor microenvironment by promoting dendritic cell maturation, enhancing NK and CD8^+^ T cell responses, and reducing immunosuppressive myeloid populations^20–24^. Like MAIT cells, iNKT cells are innately programmed for rapid effector responses and exhibit tissue-homing properties that may support their infiltration into solid tumors^4,17,21,23^.

Although both MAIT and iNKT cells are often defined by their semi-invariant TCRα chains, substantial diversity exists in their paired TCRβ chains, as well as in the overall TCR repertoire^9,18,19^. This repertoire variability may influence antigen recognition, activation thresholds, and effector functions, yet current approaches rarely capture these functional differences. Systematic characterization of MAIT TCR diversity has been limited by the lack of scalable methods that directly link TCR sequence to antigen specificity and functional response, such as how sequence variations within the MAIT TCR repertoire correlate with differences in cytokine secretion, cytotoxicity, or tissue tropism—knowledge that is critical for harnessing these cells in immunotherapy.

To address this gap, we adapted a high-throughput, single-cell functional screening platform based on hydrogel microparticles containing nanoliter-scale cavities, termed nanovials^25–29^. Our lab has previously demonstrated the utility of nanovials for the discovery of functional TCR repertoires based on selective binding to peptide major histocompatibility complex (pMHC) monomers and induced cytokine secretion^26^. In this work, we extend the nanovial platform to capture and stimulate unconventional T cells directly from human peripheral blood mononuclear cells (PBMCs) using MR1 monomers pre-loaded with 5-OP-RU and CD1d monomers pre-loaded with PBS57. This configuration allows simultaneous antigen-specific capture and activation of MAIT and iNKT cells, enabling the selective enrichment and identification of interferon gamma (IFNγ)-secreting cells without the need for pre-enrichment or complex microfluidic workflows.

Current technologies for identifying TCRs, particularly from rare or unconventional T cell subsets, often rely on peptide-MHC multimer staining and tetramer-guided sorting, followed by bulk activation assays. However, tetramer binding is not sufficient to infer function, and downstream bulk activation assays are costly, especially if applied to a large number of TCR constructs. Alternative high-throughput systems, such as microfluidic droplets or microwell arrays, have been developed for functional cytokine-based screening up front but typically require complex instrumentation, offer limited multiplexing capabilities, and do not always support downstream recovery of viable cells for transcriptomic or TCR analysis. In contrast, the nanovial platform enables single-cell resolution high-throughput screening based on functional responses—such as cytokine secretion—while maintaining cell viability and compatibility with standard flow cytometry and single-cell sequencing workflows^26,30,31^. This makes it advantageous for probing rare, functionally distinct T cells from complex human samples and for accelerating discovery of therapeutically relevant TCRs.

Here we integrated the nanovial platform with single-cell RNA and TCR sequencing using the 10x Genomics Chromium system to identify functional paired αβ TCR sequences of unconventional T cells. We used MR1-functionalized nanovials to enrich and stimulate MAIT cells and CD1d-functionalized nanovials to enrich and stimulate iNKT T cells from PBMCs of three healthy donors, sorting for sub-populations secreting IFNγ. Using secretion-encoded single-cell sequencing (SEC-seq)^25,32^, we identified and associated each TCR sequence with IFNγ secretion level and gene expression signature. We then engineered hematopoietic stem cell (HSC)-derived MAIT cells to express five of the identified MAIT TCR sequences and assessed their performance through *in vitro* and *in vivo* assays. All five sequences led to selective production of IFNγ and cell killing in co-culture assays. T cells transduced with TCR constructs from the top two functional sequences were advanced to in vivo studies and showed selective tumor infiltration and efficacy in a xenotransplant model of MR1-expressing tumor cells. Overall, the nanovial platform allows for screening of rare unconventional T cells from healthy donors based on functional properties, providing a comprehensive screening tool for discovery of rare immune cells promising for metabolism-selective cancer immunotherapies.

## Results

### Nanovials can be Functionalized to Capture and Stimulate MAIT and iNKT cells

We hypothesized that nanovials coated with MR1 or CD1d presenting their metabolite antigens would both selectively capture unconventional T cells and activate them to secrete cytokines (*Fig. 1A*). Nanovials are open cavity-containing hydrogel microparticles that can be functionalized with various proteins and antibodies. While previous work had demonstrated that T cells could be selectively loaded onto nanovials and activated by peptide-MHC monomers loaded in the nanovial cavity, we first needed to determine optimal functionalization conditions to use our nanovial platform for MR1 and CD1d. Nanovials should be coated with molecules that enable (i) isolation of MAIT and iNKT cells, (ii) stimulation of bound cells to secrete in the nanovial cavity, and (iii) capture of any secreted cytokines, such as IFNγ.

**Figure 1.**
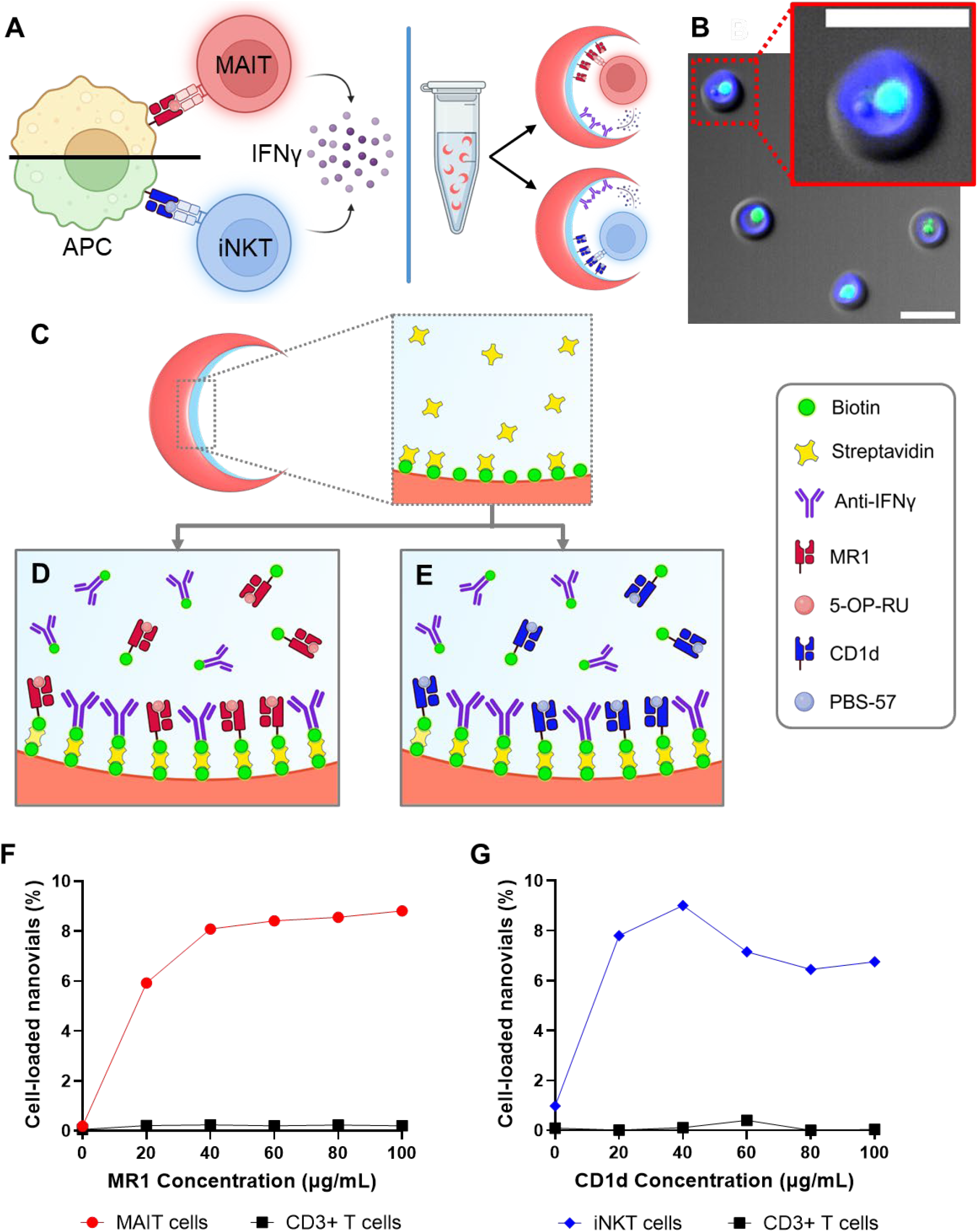
(A) Schematic of MAIT and iNKT cell engagement with their respective antigen-presenting cell (APC) leading to cytokine release. This process can be replicated in tube using nanovials conjugated with either MR1 or CD1d monomer, also leading to cytokine release. (B) Fluorescence image of MAIT cells selectively captured and activated inside nanovials to release IFNγ. Live cells are stained with calcein AM (green) and IFNγ is stained with BrilliantViolet421 (blue). Scale bar: 40μm. (C) Schematic showing functionalization of nanovial cavity using biotin-streptavidin chemistry, which are subsequently conjugated with either loaded (D) MR1 or (E) CD1d monomer and cytokine capture antibodies. (F) When loaded at a 1.6:1 cell-to-nanovial ratio, the percentage of cell-loaded nanovials for both control T cells and MAIT cells are plotted as a function of MR1 concentration on the nanovials. (G) As for **F** but using iNKT cells incubated with CD1d-functionalized nanovials.

Nanovials were formed through aqueous two-phase separation of polyethylene glycol (PEG) and gelatin in microfluidic-generated microdroplets, UV-initiated polymerization, and washing to remove the gelatin core (see methods). Following this fabrication process, lysine residues from the residual gelatin layer within each nanovial cavity were biotinylated using NHS-ester chemistry (*Fig. 1B*). This process allowed subsequent localized modification of the nanovial cavity using streptavidin and other biotinylated molecules.

For MAIT cells, as shown in *Fig. 1C*, nanovials were modified using 5-OP-RU-loaded MR1 biotinylated monomer and biotinylated anti-IFNγ capture antibodies. MR1 monomers were used to both capture and stimulate MAIT cells, while anti-IFNγ antibodies were used to detect and quantify the level of captured MAIT cell activation. Similarly, for iNKT cells, as shown in F*ig. 1D,* nanovials were modified using PBS57-loaded CD1d biotinylated monomer and biotinylated anti-IFNγ antibodies.

We first evaluated the ability of MR1 and CD1d nanovials to capture and activate MAIT and iNKT cells, respectively, from engineered T cells. Nanovials were functionalized with varying concentrations (0-100μg/ml) of either MR1 or CD1d to determine the optimal loading conditions. These nanovials were then incubated with HSC-derived unconventional T cells at 1.6:1 cell-to-nanovial ratio. A dose-response curve was established separately for each of the MAIT and iNKT cells, showing cell loading saturated to its highest levels at approximately 40µg/mL for both cell types: ∼8% of MR1-functionalized nanovials containing bound MAIT cells and ∼9% of CD1d-functionalized nanovials containing bound iNKT cells (*Fig. 1D, E*). In the absence of functionalization with MHC-like molecules, neither MAIT nor iNKT cells bind to nanovials, and control T cells lacking any unconventional TCRs failed to bind to either nanovials, confirming the specificity of the interaction

To assess the functional activation of MAIT and iNKT cells, we measured cytokine secretion after unconventional T cell loading on nanovials functionalized with either MR1 or CD1d. In a baseline control experiment, <0.2% of either MAIT or iNKT cells loaded on nanovials functionalized with anti-CD45 antibodies exhibited detectable IFNγ secretion (Fig. 2A). When nanovials were functionalized with loaded MR1 or CD1d, a significant increase in IFNγ secretion was observed. Approximately ∼17% of MAIT cells loaded on MR1-functionalized nanovials and ∼4.2% of iNKT cells loaded on CD1d-functionalized nanovials secreted IFNγ, with many cells exhibiting secretion levels more than 10-fold above background (Fig. 2B, C). This confirmed that both MR1-and CD1d-functionalized nanovials successfully induced antigen-specific cytokine release from MAIT and iNKT cells respectively.

**Figure 2.**
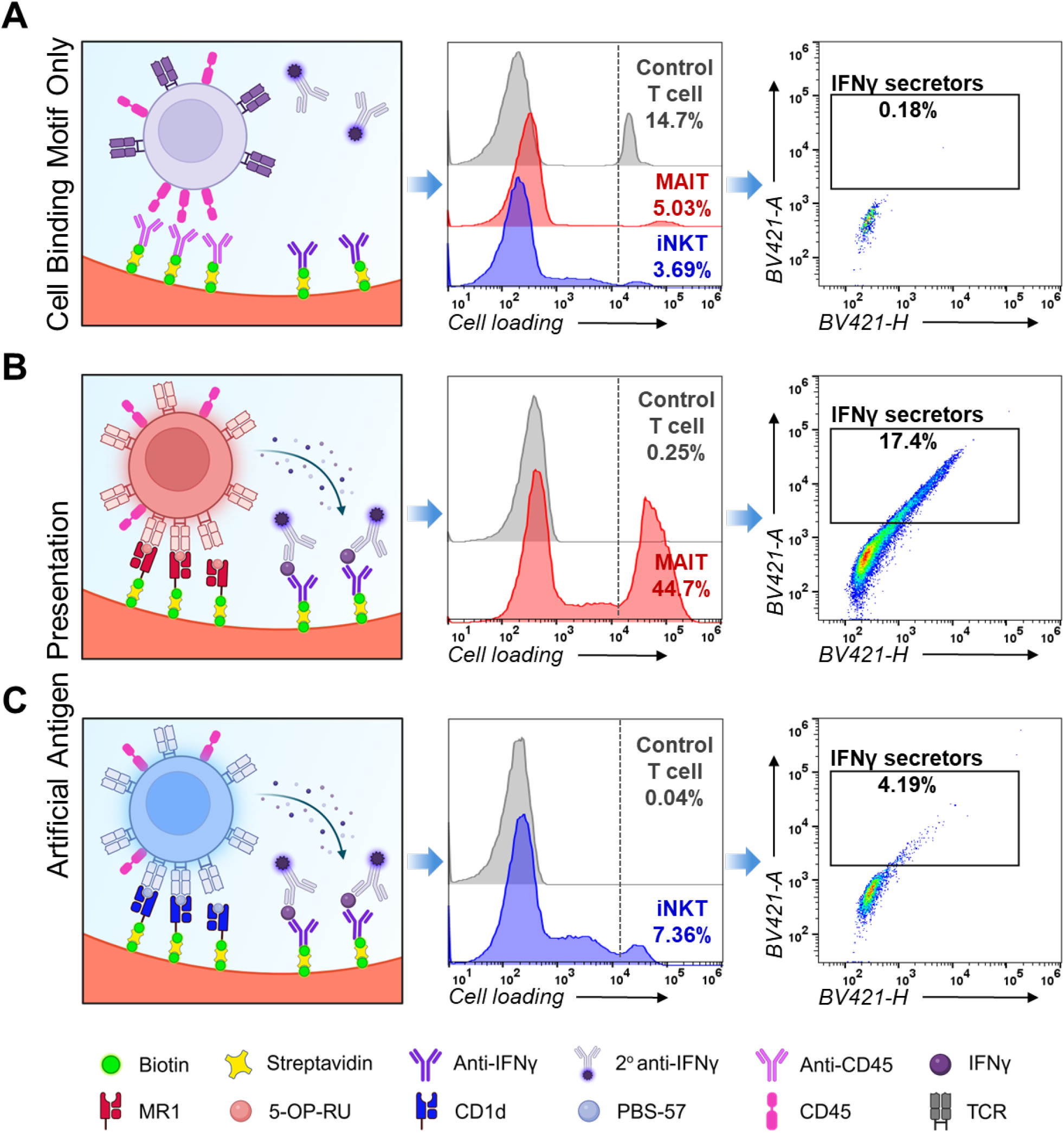
Selective binding and secretion of MAIT and iNKT cells using the nanovial platform. (A) From left to right, schematic of nanovial functionalized with anti-CD45, histogram showing non-specific capture of all types of cells, and scatter plot of bound iNKT cells showing basal level of IFNγ secretion above background in the rectangular gate. MAIT cells captured on anti-CD45 nanovials also only show basal level of IFNγ secretion (scatter plot not shown). (B) From left to right, schematic of nanovial functionalized with loaded MR1 monomer, histogram showing specific capture of MAIT cells, and scatter plot of bound MAIT cells showing increased levels of IFNγ secretion. (C) From left to right, schematic of nanovials functionalized with loaded CD1d monomer, histogram showing specific capture of iNKT cells, and scatter plot of bound iNKT cells showing increased levels of IFNγ secretion. The vertical dashed line in each histogram indicates a gate on calcein AM signal above which are live, loaded cells.

### Multiplexed Screening of PBMCs for MAIT and iNKT Cells

Next, we sought to apply the nanovial platform to identify rare MAIT and iNKT cells directly from human PBMCs. Initially, we performed experiments using fluorescently barcoded nanovials to differentiate between MR1- and CD1d-functionalized nanovials based on their distinct AF647 signals (Fig. S2). Incubating nanovials with two healthy donor PBMCs (Donor S1 and Donor S2) showed a similar percentage of cell-loaded nanovials between both donors, around 12-13%. However, flow cytometry analysis showed that the ratio of the cell-loaded nanovials comprising MAIT cells vs. iNKT cells varied between donors with ∼64% MAIT-loaded nanovials from Donor S1 as opposed to only ∼45% from donor 2 (Fig. S4). For both donors, the fraction of cells secreting IFNγ was higher upon interacting with MR1-functionalized nanovials compared to CD1d-functionalized nanovials. In Donor S1,∼2.3% of cell-loaded MR1-nanovials showed secretion signal above background compared to ∼1.4% for CD1d-nanovials (Fig. S4A). On the other hand, in Donor S2, these percentages were significantly lower at ∼0.6% for MR1-nanovials and ∼0.2% for CD1d-nanovials (Fig. S4B). Furthermore, we also showed that the frequency of IFNγ secretors in Donor S1 remained stable after three days of cell culture (Fig. S4). These experiments highlighted the donor-to-donor variability in abundance and activation level of MR1- and CD1d-specific cells.

### Single-Cell Sequencing of TCRs and Functional Markers

Using the above-mentioned combined MR1 and CD1d nanovial approach, we conducted a screen to identify MR1- and CD1d-reactive T cells from a first donor’s PBMCs (Donor 1). Reactive T cells were sorted based on the capture of secreted IFNγ levels above background and sequenced to recover matched α and β chains. We recovered a number of different clones, including two clones that possessed iNKT-specific α chains and >15 clones with MAIT-specific α chains (Table S1). Given the abundance of MAIT-specific clones, we conducted follow up studies to prioritize MAIT TCR candidates based on secretion-encoded single-cell sequencing (SEC-seq) that links IFNγ levels and gene expression directly to TCR sequences.

We conducted SEC-seq using oligo-barcodes, instead of fluorescent labels to mark the MR1- and CD1d-coated nanovials (Fig. 3). In sequential experiments, PBMCs from two different donors (Donor 2 and Donor 3) were loaded onto and incubated with MR1 and CD1d oligo-barcoded nanovials to accumulate IFNγ secretion (Fig. 3A-B), sorted for secretors (∼0.3%) (Fig. 3C) and non-secretors, labeled with anti-APC antibodies to link IFNγ levels to a separate oligonucleotide barcode (Fig. 3D), and subjected to single-cell RNA-seq (Fig. 3E). Following sequencing, we successfully recovered linked information for each single cell comprising: (i) matched αβ TCR sequences, (ii) transcriptome libraries, (iii) feature barcode counts for each nanovial type and (iv) IFNγ secretion barcode counts. We verified that cells in the IFNγ secretors gate had ∼11-fold higher mean value of IFNγ barcodes on average (Fig. S5A), providing validation that anti-APC barcode levels reflected labeling of IFNγ detection antibodies. Using the feature barcoding, we could also recover nanovials barcoded for either MR1 or CD1d, with a few double-positive and double-negative nanovials as expected (Fig. S5B). We recovered 68 MR-1 specific clones (38%), 107 CD1d-specific clones (59%), 5 double-negative clones (2.8%), and 1 double-positive clone (0.6%). Of the 68 MR-1 specific clones, we verified 100% of them to correspond to MAIT cells based on their invariant α chain. In addition, we generated a uniform manifold approximation and projection (UMAP) of the gene expression of each cell which consists of 6 different clusters, with cluster 6 showing the highest signal of barcoded IFNγ signal (Fig. S5C, D). Further analysis also showed that cells in cluster 6 exhibit higher levels of effector genes, such as granzyme B (*GZMB*) and perforin-1 (*PRF1*) (Fig. S5E, F).

**Figure 3.**
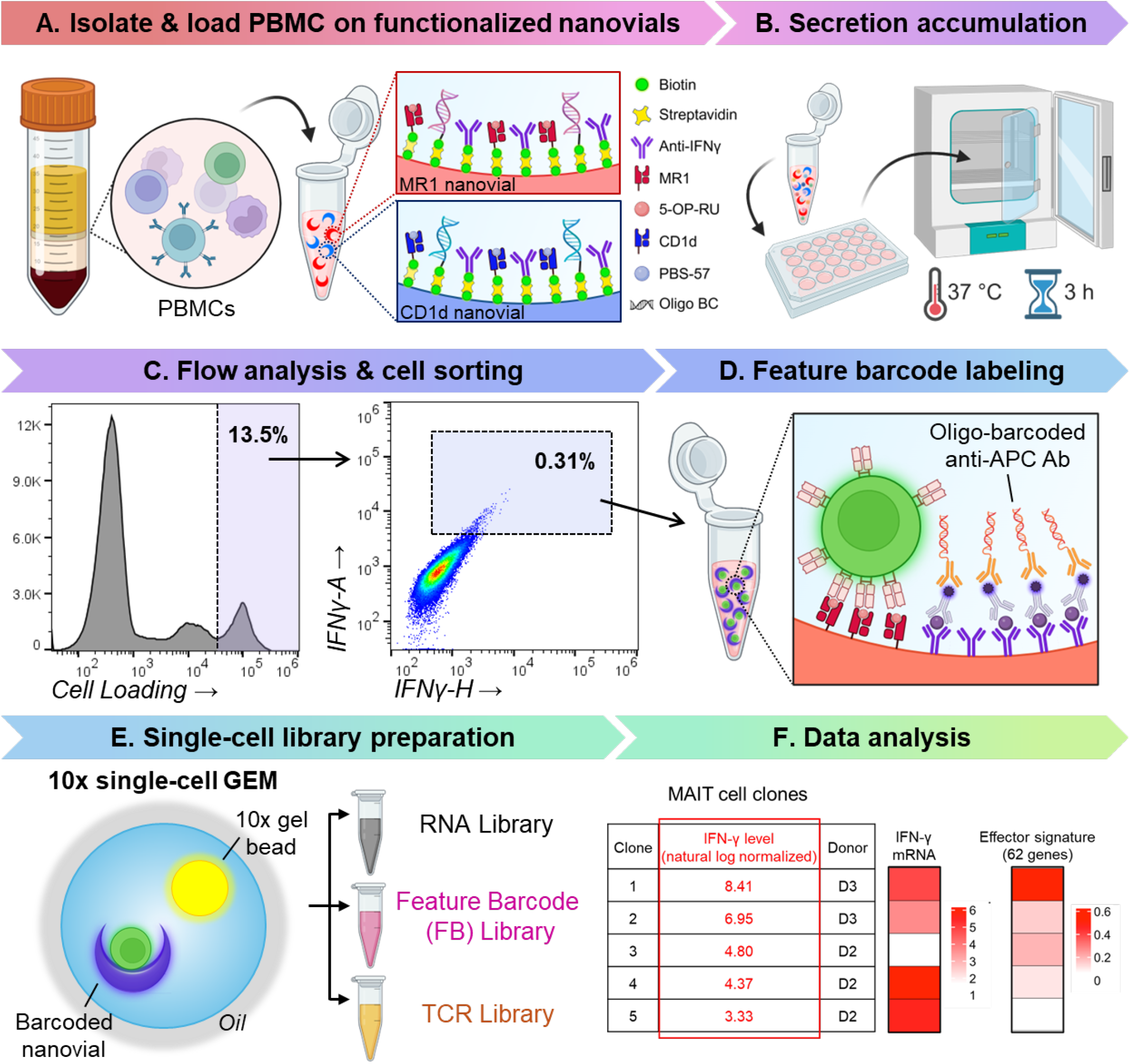
Workflow of nanovial assay to capture and analyze functional TCRs at the single-cell level. (A) Healthy donor PBMCs were isolated from whole blood and incubated with CD1d and MR1-functionalized nanovials. (B) Captured cells were activated by exposure to the nanovials to accumulate secretions for 3 hours before (C) flow analysis and FACS sorting were performed. (D) IFNγ secretor fraction was subsequently stained using oligo-barcoded anti-APC antibodies to quantify the level of IFNγ secretion during the downstream sequencing steps. (E) Single-cell cDNA libraries were prepared using the 10X Chromium system before (F) resulting data was analyzed to prioritize 5 MAIT cell clones with the highest levels of IFNγ secretion.

We selected five MAIT-specific TCRs with varying levels of IFNγ secretion (feature barcode values ranging from 27.9 to 4,490, Fig. 3F, Table S2) for further functional validation. Although *TRAJ33* was conserved across all of these clones with minimal variation in the CDR3α sequence, various paired TCRβ chains were observed (Table S2), and more highly varying CDR3β regions. The dominance of *TRAJ33* regions and increased sequence variability and length of the CDR3β regions were consistent with previously reported CDR3 sequences from human MAIT cells^33^. Interestingly, the IFNγ secretion level (Fig. S5C, D) did not correlate directly with the mRNA expression levels of IFNγ, but rather with the expression of a T cell effector gene signature (Fig. 5F). This effector gene signature included 62 genes associated with T cell activation and cytotoxicity. The T cell with the highest feature barcode for IFNγ secretion also exhibited the highest expression of these effector genes, suggesting that the level of IFNγ secretion was more closely linked to the effector state of the T cell. We further examined correlations between IFNγ secretion and additional gene signatures reflecting naïve (Tnaive), effector (Teff), terminal effector memory (Temra), effector memory (Tem), central memory (Tcm), progenitor exhausted (Tpex), and terminally exhausted (Ttex) T cell states. None of these showed a significant association with IFNγ secretion, underscoring the specific link between IFNγ production and the effector state rather than memory or exhausted phenotypes (Fig. S6).

### In Vitro Validation of MAIT TCRs

To confirm the functionality of the selected 5 MAIT TCRs, we cloned the TCRs into a lentiviral vector and transduced either human CD3-overexpressioninng 293T cells (Figure S7A, B) or primary T cells (Figure S7C). Flow cytometry confirmed successful expression of all five MAIT TCRs in the engineered human T cells. To assess their functionality, the MAIT TCR-engineered T cells were co-cultured with A375 human melanoma cells expressing MR1 (Fig. 4A). Upon stimulation with 5-OP-RU, all five TCRs induced significant IFNγ secretion, with levels higher than those observed in mock-transduced controls (Fig. 4B). Additionally, we tested the cytotoxicity of these TCR-engineered T cells against tumor cells at various effector-to-target cell ratios (10:1 to 1:1) (Fig. 4C). The engineered T cells exhibited effective killing of MR1-expressing tumor cells, with no significant increase in killing observed in mock-transduced cells or in cells incubated with tumor cells that were not treated with 5-OP-RU. Notably, TCR 1, which had the highest IFNγ secretion feature barcode and effector gene signature, demonstrated slightly enhanced tumor killing at lower effector-to-target ratios (1:1 and 2:1).

**Figure 4.**
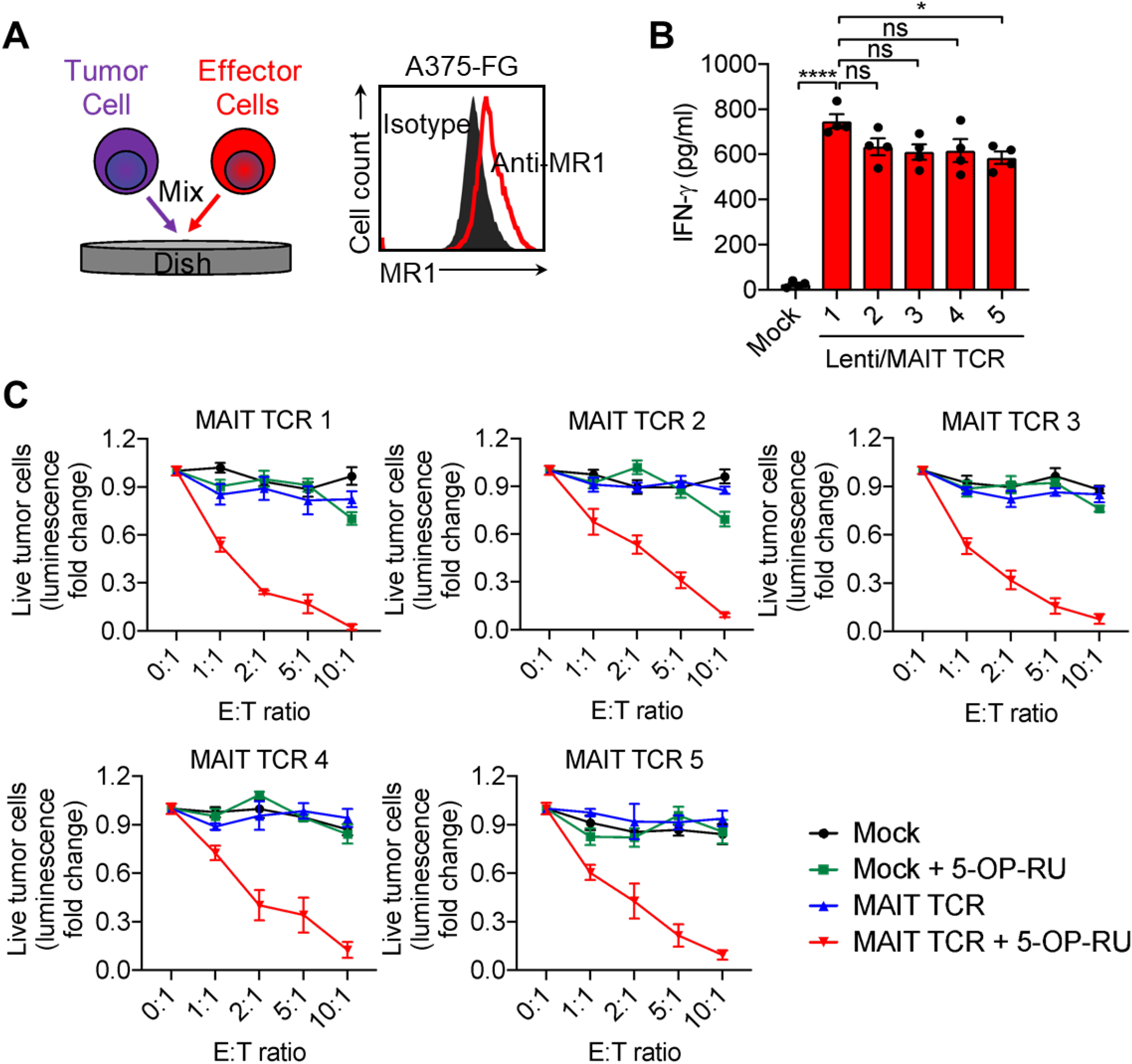
*In vitro* validation of the top 5 MAIT TCRs using *in vitro* tumor cell killing assays. (A) Experimental design to test the selected 5 MAIT TCRs using *in vitro* tumor cell killing assays, and FACS plots showing the MR1 expression on the A375-FG human melanoma cell line. A375-FG is an A375 tumor cell line overexpressing firefly luciferase and enhanced green fluorescence protein dual reporters (FG). A375-FG cells were co-cultured with therapeutic cells for 24 hours, and cytotoxicity was assessed using a bioluminescence-based assay. (B) ELISA analyses of IFNγ production by PBMC-derived MAIT TCR-engineered T cells following 24-hour co-culture with A375 tumor cells (n = 4). (C) *In vitro* tumor cell killing data for the 5 MAIT TCRs and Mock transduced controls at 24 h (n = 4).

### In Vivo Efficacy of MAIT TCR-T Cells

To evaluate the *in vivo* efficacy of the top TCR clones, we established an A375 human melanoma xenograft mouse model. On day 0, A375 cells pre-loaded with 5-OP-RU were subcutaneously implanted into NSG mice. On day 5, mice received an intravenous injection of 10 million MAIT TCR-engineered T cells (TCR 1 or TCR 2) or mock-transduced control T cells. Tumor volumes were monitored every 5 days, and animals were sacrificed on day 30 for endpoint analyses (Fig. 5A and Fig. S8A). Both TCR 1 and TCR 2 significantly suppressed tumor growth, as demonstrated by reduced tumor volume and weight compared to mock-transduced controls (Fig. 5B, C and S8A). This tumor control persisted throughout the 30-day study period. Flow cytometry analysis of tumor-infiltrating human T cells showed that 10-15% of the cells were human T cells, and more than 90% of these infiltrating cells stained positive for the engineered TCR (Vα7.2) in both TCR 1 and TCR 2 groups. Notably, TCR-T cells also exhibited significantly higher levels of CD69 activation marker staining and intracellular IFNγ compared to mock-transduced cells, indicating active T cell engagement within the solid tumor microenvironment (Fig. 5D-G, Fig. S8B, C). Collectively, these findings validate the antitumor functionality of MAIT TCRs identified through the nanovial-based selection platform and underscore the therapeutic potential of MAIT TCR-engineered T cells for solid tumor immunotherapy.

**Figure 5.**
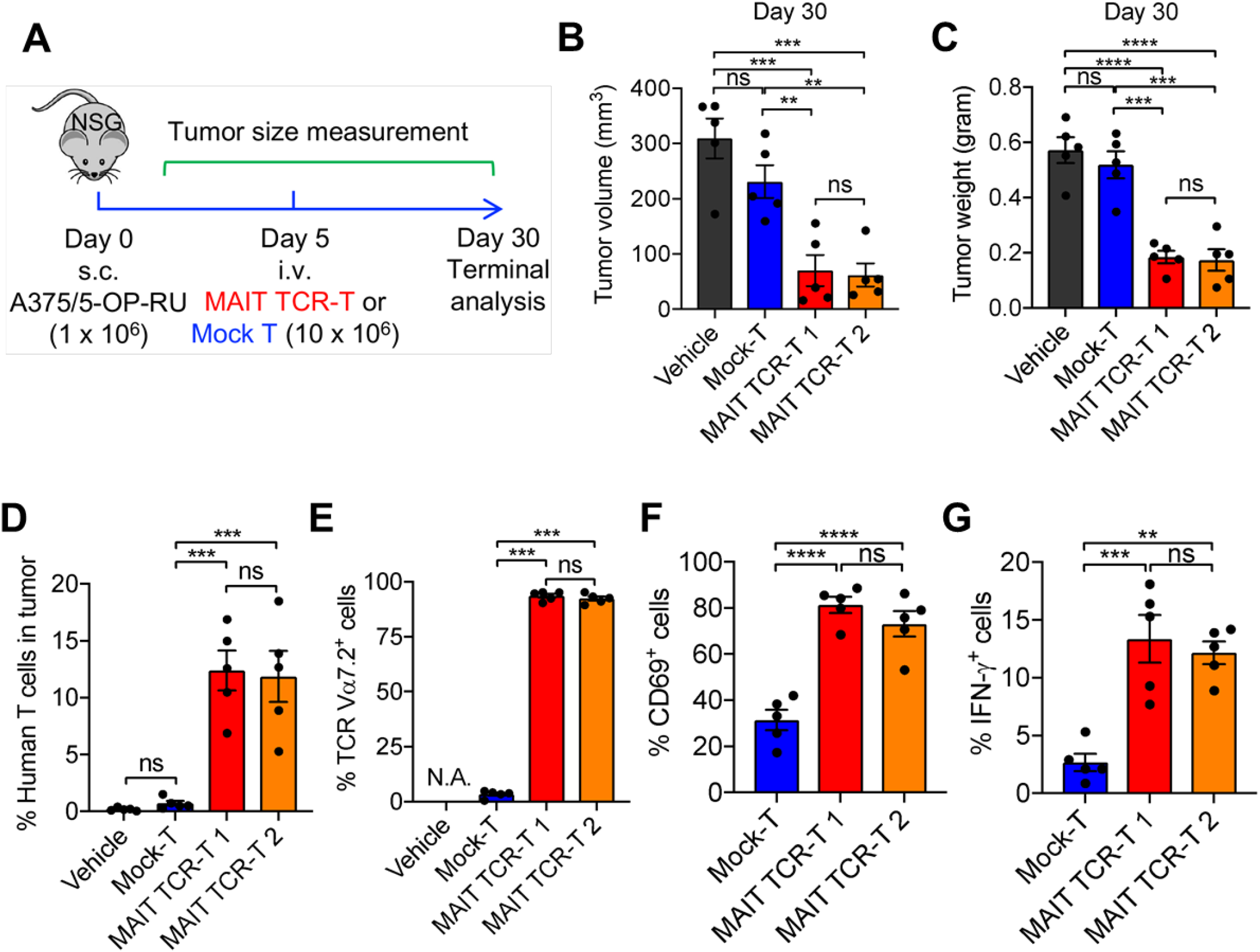
*In vivo* validation using the top 2 MAIT TCRs using an A375 human melanoma xenograft mouse model. (A) Experimental design. s.c. subcutaneous; i.v., intravenous. (B) Tumor volume measurement on day 30 (n = 5). (C) Tumor weight measurement on day 30 (n = 5). (D) FACS analyses of human T cell infiltration into the solid tumors (n = 5). (E) FACS analyses of MAIT TCR-engineered T cell proportion in the tumor-infiltrating T cells (n = 5). (F) FACS analyses of T cell activation marker CD69 expression on the tumor-infiltrating T cells (n = 5). (G) FACS analyses of effector cytokine IFNγ in the tumor-infiltrating T cells (n = 5). Statistical analysis was done using one-way ANOVA; ns, not significant; *p<0.05; **p<0.01; ***p<0.001;****p<0.0001.

## Discussion

In this study, we developed and validated a nanovial-based single-cell functional screening platform for the discovery of TCRs from unconventional T cells, with a focus on MAIT and iNKT subsets. By functionalizing nanovials with MR1 and CD1d monomers loaded with known non-peptide antigens, we were able to selectively capture and stimulate antigen-specific cells from both engineered cell products and primary PBMCs, enriching for functional cells based on IFNγ secretion. Importantly, our platform enabled high-throughput identification of functional cells while remaining compatible with downstream flow cytometry and single-cell RNA and TCR sequencing. This approach overcomes key limitations of current methods, which often rely on tetramer-guided sorting and bulk functional validation assays that fail to capture functional heterogeneity or require complex, low-throughput workflows.

In the multiplex nanovial studies, we observed donor-to-donor variability in both the frequency and functional activity of MAIT and iNKT cells, consistent with prior reports suggesting dynamic regulation of these cell types in circulation by microbial exposure, cytokine environment, and host genetics. Notably, our assay does not pre-activate or condition cells prior to the 3-hour stimulation on nanovials, using only presented antigen (signal 1), without additional signaling implicated in long-term T cell responses, which may also yield a smaller set of already primed cells. Integrating SEC-seq into our assay, we were able to associate each recovered TCR with its corresponding cytokine secretion profile and transcriptomic signature. This enabled us to identify and rank candidate MAIT TCRs based on their functional activity and effector state. Further validation of these TCRs in engineered primary T cells also confirmed their effectiveness both in vitro and in vivo. The use of fluorescent- and oligonucleotide-barcoded nanovials for multiplexed screening from a single sample demonstrated the potential of nanovial platform to simultaneously profile multiple unconventional T cell subsets.

The nanovial platform can be further expanded for a broad range of unconventional T cell subsets, beyond MAIT and iNKT cells. Since nanovials can be modularly functionalized with a variety of non-classical MHC-like molecules—including MR1, CD1d, and potentially CD1b, CD1c, or even BTN or HLA-E—this system is readily adaptable to diverse metabolite- or lipid-reactive T cells. This versatility expands the potential for systematic, function-first screening across unconventional T cell populations, many of which remain poorly characterized yet display tumor reactivity and favorable immunotherapeutic potential.

In addition, as discussed, the nanovial platform supports antigen-specific activation without requiring co-stimulatory signal 2, a critical feature for evaluating unconventional T cells whose activation is often TCR-dominant or cytokine-enhanced. The absence of a required co-stimulatory signal avoids artificial amplification of low-affinity responses and ensures that secretion and transcriptomic profiles reflect direct antigen recognition. Our data and previous study also validate that both peptide–MHC complexes and metabolite-presenting molecules like MR1 and CD1d remain functionally active when immobilized within the nanovial cavity, maintaining proper antigen presentation for TCR activation.

Our system provides an experimental framework for future studies coupling nanovial-based capture with antigen discovery techniques such as MR1 ligand loading from tumor lysates and identification by mass spectrometry. By integrating these pipelines, we envision the ability to not only identify reactive TCRs but also elucidate their cognate antigens, thus advancing the design of targeted, metabolism-informed immunotherapies. In addition, expanding this platform to profile TCRs from tumor-infiltrating unconventional T cells, or from cancer patients, may also reveal novel receptors with enhanced tumor selectivity or resistance to immunosuppressive signaling.

The nanovial platform’s broad compatibility, independence from co-stimulatory signals, and robust support for antigen presentation and functional readouts position it as a powerful tool for unconventional T cell profiling and TCR discovery. These attributes may help unlock novel therapeutic modalities rooted in the unique biology of T cells that recognize non-peptidic antigens and operate outside the constraints of HLA polymorphism.

## Methods

### Nanovial Fabrication

Nanovials were manufactured using a PDMS-based flow-focusing microfluidic device, following previously established protocols^29^. In this setup, three precursor solutions—polyethylene glycol (PEG), gelatin, and oil— were co-flowed into the droplet generator. The PEG phase consisted of 27.5% (w/v) 4-arm, 5 kDa PEG-acrylate (Advanced BioChemicals) and 4% (w/v) lithium phenyl-2,4,6-trimethylbenzoylphosphinate (Sigma-Aldrich) dissolved in 1X phosphate-buffered saline (PBS; ThermoFisher Scientific). The gelatin solution was prepared by dissolving 20% (w/v) cold-water fish skin gelatin (Sigma-Aldrich) in sterile MilliQ water. The oil phase was composed of 0.5% (v/v) Pico-Surf (Sphere Fluidics) in Novec™ 7500 Engineered Fluid (3M). These phases were introduced into the device at flow rates of 1, 1, and 10 μL/min for PEG, gelatin, and oil respectively. After droplet formation and phase separation, UV light was applied to crosslink the PEG network, forming nanovial particles. Emulsions containing the crosslinked particles were demulsified using 20% (v/v) perfluoro-1-octanol (PFO; Sigma-Aldrich) in Novec™ 7500. Residual oil was removed by sequential hexane washes followed by centrifugation, repeated twice. To eliminate the remaining hexane, the particles were washed with sterile 70% ethanol, centrifuged, and aspirated. Nanovials were then functionalized by overnight incubation at 4°C with 10 mM Sulfo-NHS-Biotin (APExBIO). Final sterilization was performed using 70% ethanol, and nanovials were stored at 4°C in Particle Wash Buffer (1X Dulbecco’s PBS (Gibco™), 0.05% Pluronic F-127 (Sigma), 0.5% Bovine Serum Albumin (Sigma-Aldrich), and 1X Antibiotic-Antimycotic (Gibco™)).

### Nanovial Functionalization

#### Streptavidin Conjugation

Nanovials were diluted to approximately 37,000 particles/mL in Particle Wash Buffer and incubated with an equal volume of streptavidin solution (200μg/mL; ThermoFisher Scientific) for 30 minutes at room temperature on a tube rotator. Following incubation, excess streptavidin was removed by brief centrifugation using a benchtop minifuge, and the supernatant was carefully aspirated. To ensure complete removal of unbound reagent, nanovials were resuspended in fresh Particle Wash Buffer, centrifuged again, and the supernatant was discarded. This washing procedure was repeated two additional times.

#### MHC-like Monomers and Antibody Conjugation

Functionalization of control anti-CD45 nanovials were done by incubating 37,000 particles/mL nanovials with 20μg/mL (130nM) of each biotinylated anti-CD45 (BioLegend) and biotinylated anti-IFNγ (R&D Systems). For antigen-specific stimulation, MR1 and CD1d nanovials were similarly prepared at the same particle concentration using 40μg/mL of biotinylated MR1 (pre-loaded with 5-OP-RU) or CD1d (pre-loaded with PBS57) monomers, along with 20μg/mL biotinylated anti-IFNγ (R&D Systems). In monomer titration experiments, only the concentration of the MR1 or CD1d monomer was varied, while the concentration of anti-IFNγ remained constant. Functionalization was done overnight at 4°C, followed by removal of unbound reagents and two washes.

### Generation of pooled MR1- and CD1d-functionalized nanovials for multiplex assay

For the proof-of-concept multiplex experiment, nanovials to be functionalized with CD1d were first incubated with fluorescently labeled streptavidin (SA-AF647) at 1:100 ratio with regular streptavidin (Fig. S3) before removal of excess of reagents and subsequent washes. After the functionalization step, MR1- and AF647-labeled CD1d-nanovials were pooled together at 1:1 ratio use.

TotalSeq™ Oligomer-conjugated streptavidin was used as barcodes, instead of SA-AF647, to differentiate between MR1 and CD1d nanovials for downstream single cell analysis. MR1 nanovials were labeled with TotalSeq™-C0971 Streptavidin (BioLegend), while CD1d nanovials were labeled with TotalSeq™-C0975 Streptavidin. Similar to above, after functionalization and barcoding, MR1- and CD1d-nanovials were pooled together at 1:1 ratio before use.

### Isolation of IFNγ-secreting MAIT and iNKT cells in nanovials

Healthy donor PBMCs from UCLA CFAR/Virology core were loaded into pooled MR1 and CD1d nanovials at 5:1 cell-to-nanovial ratio for 1.5 hours with constant gentle agitation at 37°C, followed by removal of unbound cells using cell strainer. Bound cells were then stimulated by the nanovials for 3 hours at 37°C. After, samples were stained with calcein AM solution (1:5000 dilution in Particle Wash Buffer) and APC/Cy7-conjugated IFNγ detection antibody (5uL per ∼200,000 nanovials). Antibody staining was performed for 30 minutes at 37°C. Samples were washed with Particle Wash Buffer twice to remove excess fluorescent dyes. Flow cytometry analysis and FACS-based sorting was done using SONY SH800. For SEC-seq experiments, sorted cells were then spun down and labeled with TotalSeq™-C0987 anti-APC Antibody (BioLegend) for 30 minutes at 4°C. Sorted samples were then washed with and resuspended in 0.04% BSA in PBS.

### Single cell sequencing and identification of MAIT TCRs

Freshly sorted samples of PBMC-derived MAIT cells in nanovials were immediately transferred to the UCLA TCGB Core for single-cell RNA sequencing. The number of cell-loaded nanovials loaded onto the Chromium Controller (10x Genomics) was based on the number of events sorted by FACS. Gene-expression, T-cell receptor (TCR) V(D)J, and feature-barcoding libraries were prepared with the Chromium Next GEM Single Cell 5’ v2 Gene Expression Kit, the Human T-cell V(D)J Enrichment Kit, and the Feature Barcoding Kit following the manufacturers’ protocols. Library quality was assessed on a 4200 TapeStation with D1000 ScreenTape (Agilent) and quantified by Qubit. Pooled libraries were sequenced on an Illumina NovaSeq 6000 S4 flow cell (2 × 100 bp).

FASTQ files were processed with Cell Ranger Count v7.2.0 (gene-expression and feature-barcode) and with Cell Ranger V(D)J v7.2.0 (TCR sequencing) against GRCh38 reference genome, yielding gene-expression, feature-barcode, and paired α/β-chain contig matrices.

Analyses were performed in Seurat v4. Gene-expression data and feature-barcode UMI counts were log-normalized and scaled. Clonotypes were defined by identical paired CDR3α/CDR3β nucleotide sequences. For gene signatures, per-cell module scores were calculated with AddModuleScore^34^. The five dominant clonotypes were selected, and their IFNγ secretion levels (feature-barcode counts) and module scores were compared by violin plots and Wilcoxon rank-sum tests.

### Primary human T cell culture for generation of engineered MAIT cells

PBMCs from healthy donors were used to expand T cells for additional engineering. T cell activation was achieved using one of two methods: (1) stimulation with Dynabead™ Human T-Activator CD3/CD28 (Thermo Fisher Scientific, Cat. No. 11131D) according to the manufacturer’s instructions, or (2) plate-bound activation. For the latter, non-treated 24-well tissue culture plates (Corning, Cat. No. 3738) were coated with Ultra-LEAF*TM* purified anti-human CD3 antibody (Clone OKT3, BioLegend, cat. no. 317325) at 1 µg/ml (500 µl/well) for 2 hours at room temperature or overnight at 4 °C. PBMCs were then resuspended in C10 medium supplemented with 1 µg/ml Ultra-LEAFTM purified anti-human CD28 antibody (Clone CD28.2, BioLegend, cat. no. 302933) and 30 ng/ml IL-2, and seeded into the pre-coated plates at a density of 1 × 10^6^ cells/ml (1 ml/well). Following activation, cells were maintained in C10 medium supplemented with 20 ng/ml IL-2 and cultured for 2-3 weeks.

### Generation and culture of engineered MAIT cells

A parental lentivector, pMNDW, was utilized to construct the lentiviral vectors employed in this study. The 2A sequences derived from foot-and-mouth disease virus (F2A) were used to link the inserted genes to achieve co-expression. The Lenti/MAIT vector was generated by inserting into the pMNDW parental backbone a synthetic bicistronic gene encoding human MAIT TCRα-F2A-MAIT TCRβ. After 2 days of plate-bound activation, T cells were transduced with Lenti/MAIT viruses for a period of 24 hours. The engineered T cells were expanded for about 2 weeks in C10 medium and then cryopreserved for future applications.

### Generation and culture of target tumor cells

Human melanoma cell line A375 (cat. no. CRL-1619) was purchased from the American Type Culture Collection (ATCC). To establish stable tumor cell lines that overexpress firefly luciferase and green fluorescent protein dual reporters (FG), the parental tumor cell lines were transduced with lentiviral vectors carrying the specific genes of interest (i.e., Lenti/FG). 72h after lentiviral transduction, the cells underwent flow cytometry sorting to isolate the genetically modified cells (as identified as GFP^+^ cells) necessary for creating stable cell lines.

### In vitro tumor cell killing assay

Human tumor cells (e.g., A375-FG; 1 × 10^4^ cells per well in 96-well plate) were co-cultured with the indicated therapeutic cells (i.e., mock T or MAIT TCR-engineered T cells) in Corning 96-well clear bottom black plates for 24 hours in C10 medium. The effector cell to target cell (E:T) ratio is indicated in the figure legends. At the end of culture, viable tumor cells were quantified by adding D-luciferin (150 μg/ml; Fisher Scientific, cat. no. 50-209-8110) to cell cultures, followed by the measurement of luciferase activity using an Infinite M1000 microplate reader (Tecan).

### In vivo A375 human melanoma xenograft mouse model

Experimental design is shown in Figure 5A. Briefly, on Day 0, NSG mice received a subcutaneous (s.c.) injection of A375 human melanoma cells (1 × 10^6^ cells per mouse). On Day 5, the experimental mice received an intravenous (i.v.) injection of mock T or MAIT TCR-engineered T cells (10 × 10^6^ cells in 100 μl PBS per mouse).

Over the experiment, the tumor load was measured using tumor size measurements. At the study endpoint, experimental mice were euthanized, and the tumor-infiltrating T cells were analyzed using flow cytometry.

Fluorochrome-conjugated antibodies specific for human CD3 (Clone HIT3a, Pacific Blue, PE, or PE-Cy7-conjugated, 1:500, cat. no. 300330, 300308, or 300316), CD45 (Clone HI30, PerCP, FITC, or Pacific Blue-conjugated, 1:500, cat. no. 982318, 982316, or 982306), CD69 (Clone FN50, PE-Cy7 or PerCP-conjugated, 1:50, cat. no. 310912 or 310928), IFNγ (Clone B27, PE-Cy7-conjugated, 1:50, cat. no. 506518), TCR Vα7.2 (Clone 3C10, PE-conjugated, 1:50, cat. no. 351706) were purchased from BioLegend. Fixable Viability Dye eFluor506 (e506, 1:500, cat. no. 65-0866-14) was purchased from Affymetrix eBioscience; mouse Fc Block (anti-mouse CD16/32, cat. no. 553141) was purchased from BD Biosciences; and human Fc Receptor Blocking Solution (TrueStain FcX) was purchased from BioLegend (cat. no. 422302). In our study, we note the use of antibodies with identical clones but differing conjugated fluorochromes, with one typical antibody listed herein. All FACS staining was performed following manufacturers’ provided protocols. Appropriate isotype staining controls were used for all staining procedures. Stained cells were analyzed using a MACSQuant Analyzer 10 flow cytometer (Miltenyi Biotech), following the manufacturer’s instructions. FlowJo software version 9 (BD Biosciences) was used for data analysis.

## Acknowledgements

We would like to acknowledge the UCLA Jonsson Comprehensive Cancer Center for their flow cytometry services and UCLA/CFAR Virology Core Laboratory for providing primary human PBMCs. Y.-R.L. is a postdoctoral fellow supported by a UCLA MIMG M. John Pickett Post-Doctoral Fellow Award, a CIRM-BSCRC Postdoctoral Fellowship, a UCLA Sydney Finegold Postdoctoral Award, a UCLA Chancellor’s Award for Postdoctoral Research, and a UCLA Goodman-Luskin Microbiome Center Collaborative Research Fellowship Award. Y.Z. is a pre-doctoral fellow supported by a a Whitcome pre-doctoral fellowship in molecular biology.

This project was supported by grant 2023-332384 from the Chan Zuckerberg Initiative Donor Advised Fund (CZI DAF), an advised fund of the Silicon Valley Community Foundation. We thank the NIH Tetramer Core Facility (NIH Contract 75N93020D00005 and RRID:SCR_026557) for providing both MR1 and CD1d monomers used in this study. Figures were created in https://BioRender.com.

