## Supplementary Information for "A Nanovial-Based Platform for Functional Discovery of Antigen-Reactive TCRs from Unconventional T Cells"

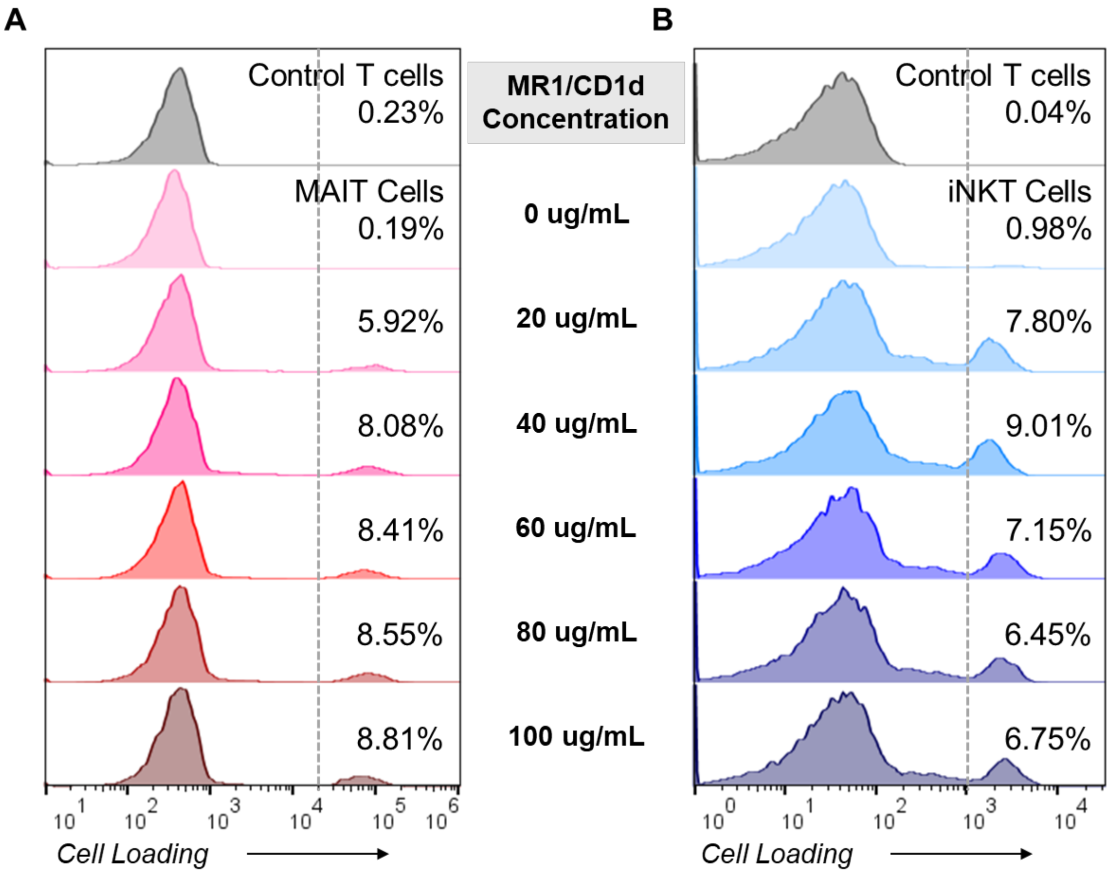


**Figure S1.** (A) Flow cytometry histograms showing percentage of MAIT cell-loaded nanovials at different concentrations of MR1 functionalization of nanovials. (B) As for **A** with iNKT cell-loaded nanovials across different concentrations of CD1d.

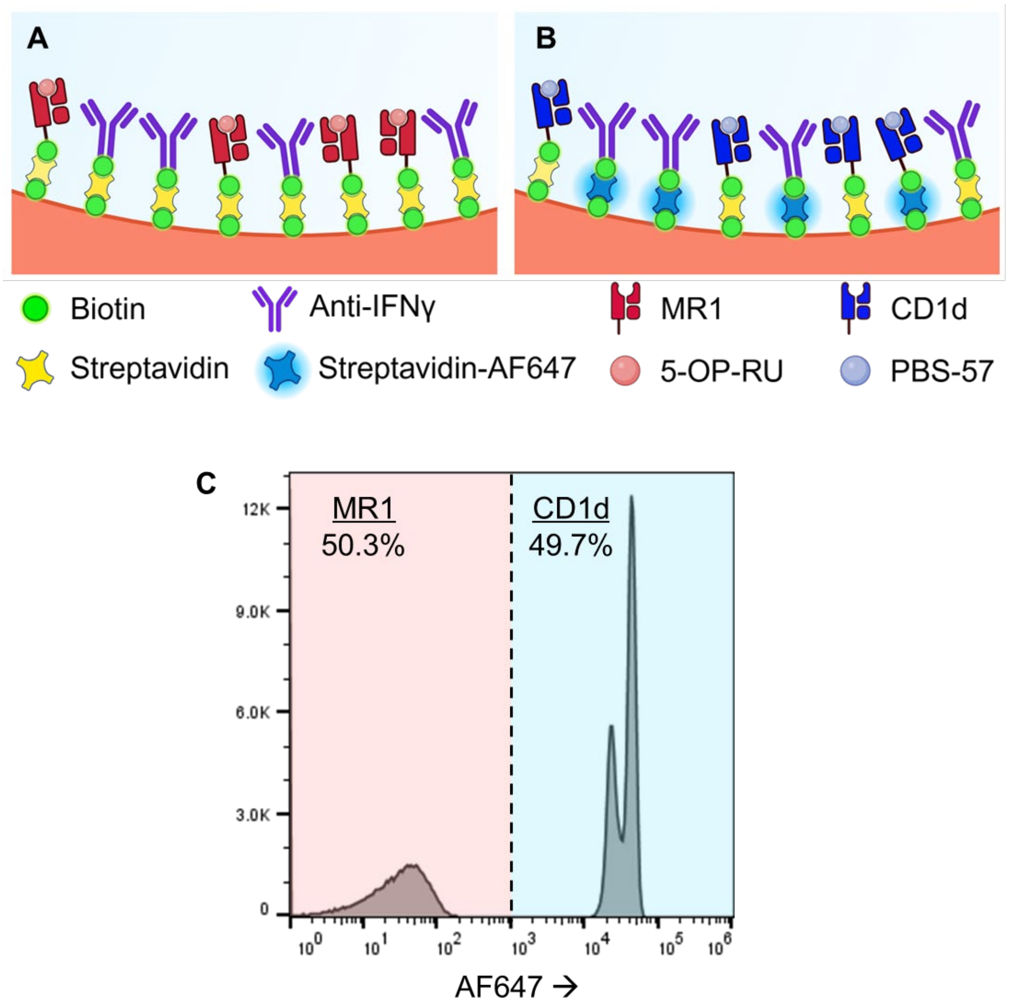


**Figure S2.** Schematics showing (A) nanovials functionalized with MR1 and anti-IFNγ antibodies and (B) nanovials first incubated with a mix of streptavidin and streptavidin-AF647, before subsequent functionalization with CD1d and anti-IFNγ antibodies. The two types of nanovials were pooled together at 1:1 ratio after being functionalized. (C) Flow cytometry histogram plot showing higher AF647 signal on MR1 nanovials with the ability to gate MR1 and CD1d nanovials based on their fluorescence.

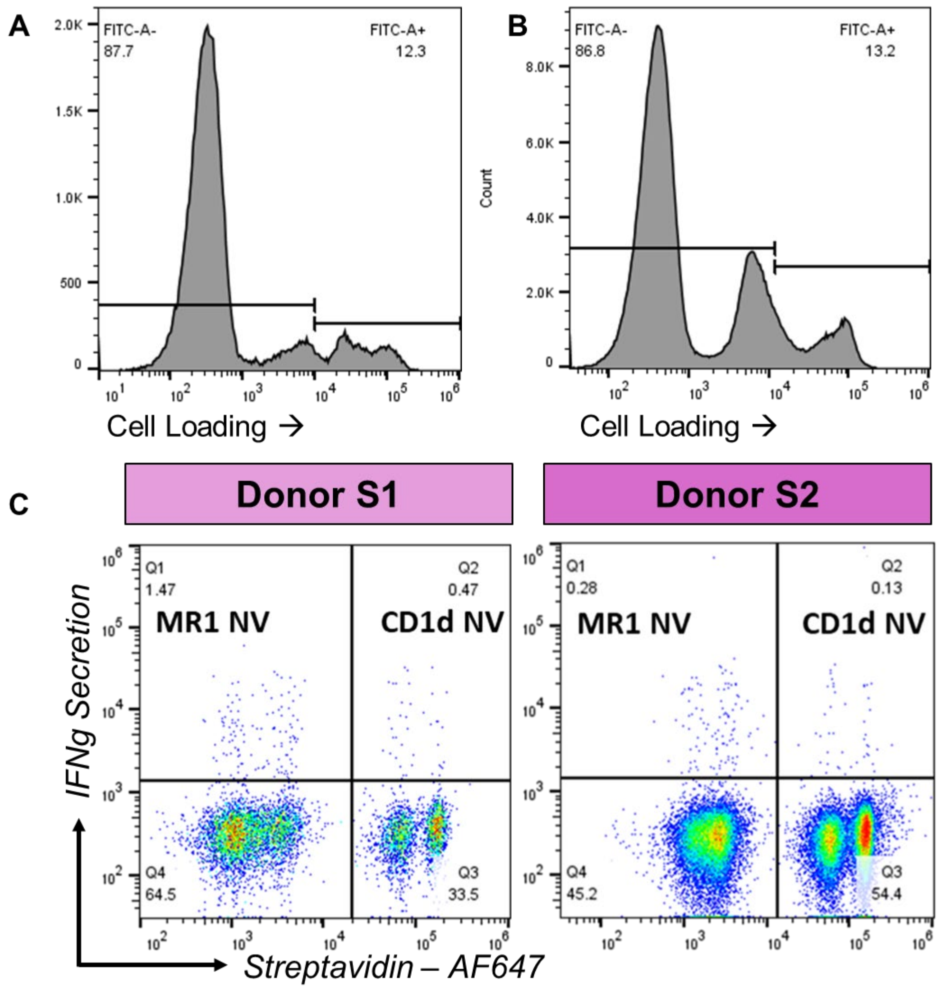


**Figure S3.** Histograms showing the distribution of calcein AM staining of cell-loaded nanovials for (A) Donor S1 and (B) Donor S2. High calcein AM components of the distributions correspond to viable loaded cells. Unloaded nanovials appear as the components of the distributions with a low fluorescence peak. (C) Flow plots showing distributions of IFNγ secretion for MR1 (low AF647)- and CD1d (high AF647)-specific cells that bound to nanovials for both donors.

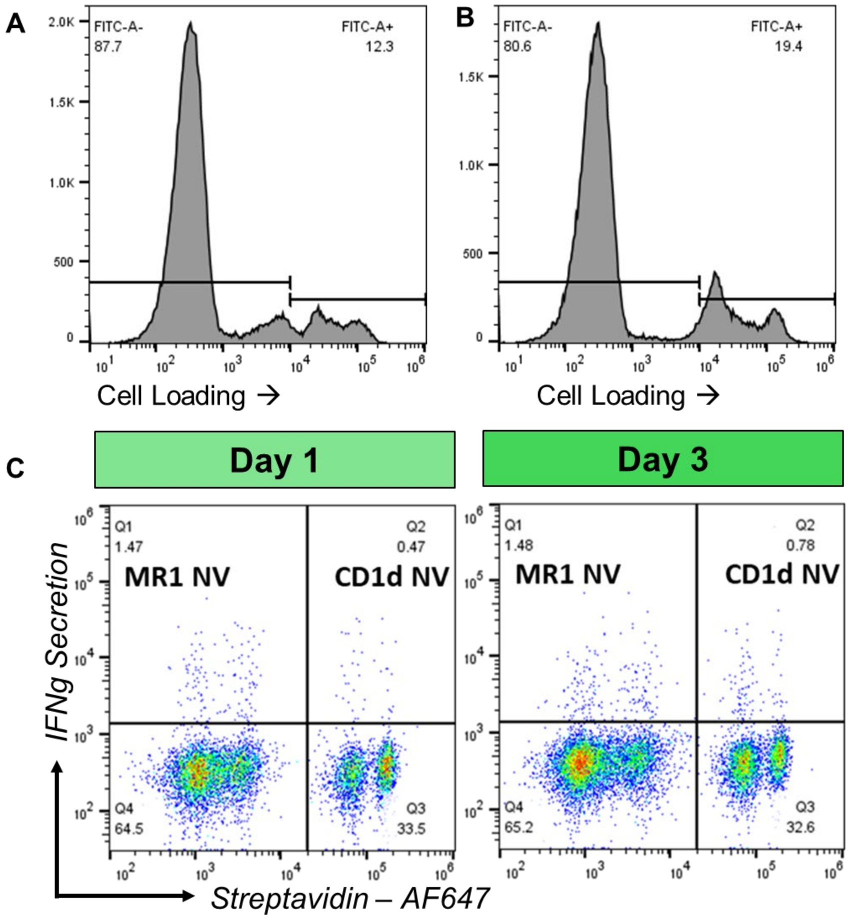


**Figure S4.** Histograms showing the distribution of calcein AM staining of cell-loaded nanovials for (A) day 1 and (B) day 3 following Donor S1 PBMC isolation. High calcein AM components of the distributions correspond to viable loaded cells. Unloaded nanovials appear as the components of the distributions with a low fluorescence peak. (C) Flow plots showing distributions of IFNγ secretion for MR1 (low AF647)- and CD1d (high AF647)-specific cells that bound to nanovials for both donors.

**Table 1.** Sequencing results for Donor 1 showing iNKT (TRAV10,TRAJ18) and MAIT (TRAV1-2,TRAJ33) specific α-chain genes recovered.

| Clonotype ID | CDR3α | TRAV | TRAJ | TRAC | CDR3β | TRBV | TRBJ | TRBC | Identification |
|---|---|---|---|---|---|---|---|---|---|
| 1 | CLQSARLMF<br>CVVSDRGSTLGRLYF | TRAV4<br>TRAV10 | TRAJ31<br>TRAJ18 | TRAC<br>TRAC | CASSERNGSTDTQYF | TRBV25-1 | TRBJ2-3 | TRBC2 | **Mixed** |
| 2 | CVVSDRGSTLGRLYF | TRAV10 | TRAJ18 | TRAC | CASSSRGPDGQFF | TRBV25-1 | TRBJ2-1 | TRBC2 | **iNKT** |
| 3 | CAPMDSNYQLIW | TRAV1-2 | TRAJ33 | TRAC | CASSEESSGSYEQYF | TRBV6-1 | TRBJ2-7 | TRBC2 | **MAIT** |
| 4 | CAVVDSNYQLIW | TRAV1-2 | TRAJ33 | TRAC | CSARDSTSGSEMETQYF | TRBV20-1 | TRBJ2-5 | TRBC2 | **MAIT** |
| 5 | CAVMDSNYQLIW | TRAV1-2 | TRAJ33 | TRAC | CSANIDRATEAFF | TRBV20-1 | TRBJ1-1 | TRBC1 | **MAIT** |
| 6 | CAVLDSNYQLIW | TRAV1-2 | TRAJ33 | TRAC | CSADRGLLSDGYTF | TRBV20-1 | TRBJ1-2 | TRBC1 | **MAIT** |
| 7 | CAVLDSNYQLIW | TRAV1-2 | TRAJ33 | TRAC | CSARNQIGSDTEAFF | TRBV20-1 | TRBJ1-1 | TRBC1 | **MAIT** |
| 8 | CAAYVSGNTPLVF | TRAV29/DV5 | TRAJ29 | TRAC | CASSLGGSYGYTF | TRBV5-6 | TRBJ1-2 | TRBC1 | Unknown |
| 9 | CAVRDSNYQLIW | TRAV1-2 | TRAJ33 | TRAC | CASSYDVSTDTQYF | TRBV6-2 | TRBJ2-3 | TRBC2 | **MAIT** |
| 10 | CAVLDSNYQLIW | TRAV1-2 | TRAJ33 | TRAC | CATSPGGDAGELFF | TRBV24-1 | TRBJ2-2 | TRBC2 | **MAIT** |
| 11 | CAVQEGGGGSYIPTF | TRAV20 | TRAJ6 | TRAC | CASNSGGQEKLFF<br>CASSEGTSGSYEQYF | TRBV2<br>TRBV6-1 | TRBJ1-4<br>TRBJ2-7 | TRBC1<br>TRBC2 | **Mixed** |
| 12 | CAVMDSNYQLIW | TRAV1-2 | TRAJ33 | TRAC | CASSRTGGNTGELFF | TRBV6-4 | TRBJ2-2 | TRBC2 | **MAIT** |
| 13 | CAVMDSNYQLIW | TRAV1-2 | TRAJ33 | TRAC | CASSYRLAGAYEQYF | TRBV6-5 | TRBJ2-7 | TRBC2 | **MAIT** |
| 14 | CAVMDSNYQLIW | TRAV1-2 | TRAJ33 | TRAC | CATIIGRGENSPLHF | TRBV6-1 | TRBJ1-6 | TRBC1 | **MAIT** |
| 15 | CAVMDSNYQLIW | TRAV1-2 | TRAJ33 | TRAC | CASSEAGGPTYEQYF | TRBV6-1 | TRBJ2-7 | TRBC2 | **MAIT** |
| 16 | CAVMDSNYQLIW | TRAV1-2 | TRAJ33 | TRAC | CASSEGTSGYNEQFF | TRBV6-1 | TRBJ2-1 | TRBC2 | **MAIT** |
| 17 | CAVEDQGYNQGGKLIF<br>CAVMDSNYQLIW | TRAV2<br>TRAV1-2 | TRAJ23<br>TRAJ33 | TRAC<br>TRAC | CASSEVAGGTDTQYF | TRBV6-1 | TRBJ2-3 | TRBC2 | **Mixed** |
| 18 | CAALDSNYQLIW | TRAV1-2 | TRAJ33 | TRAC | CASTPRTERTDTQYF | TRBV28 | TRBJ2-3 | TRBC2 | **MAIT** |
| 19 | CAVLDSNYQLIW | TRAV1-2 | TRAJ33 | TRAC | CATSPDRGGTDTQYF | TRBV15 | TRBJ2-3 | TRBC2 | **MAIT** |
| 20 | CAVTDSNYQLIW | TRAV1-2 | TRAJ33 | TRAC | CASSDSTSGTTLPQFF | TRBV6-4 | TRBJ2-1 | TRBC2 | **MAIT** |
| 21 | CAVRDSDYKLSF | TRAV1-2 | TRAJ20 | TRAC | CASSDSAGTGTDTQYF | TRBV6-4 | TRBJ2-3 | TRBC2 | **MAIT** |
| 22 | CAVMDSNYQLIW | TRAV1-2 | TRAJ33 | TRAC | CATSRVAGGGTDTQYF | TRBV15 | TRBJ2-3 | TRBC2 | **MAIT** |
| 23 | CIVRSNFGNEKLTF | TRAV26-1 | TRAJ48 | TRAC | CASSPQQGAIYGYTF | TRBV12-3 | TRBJ1-2 | TRBC1 | Unknown |
| 24 | CAVLDSNYQLIW | TRAV1-2 | TRAJ33 | TRAC | CASSEEVGGTGGNSDTQYF | TRBV6-1 | TRBJ2-3 | TRBC2 | **MAIT** |
| 25 | CVVSDRGSTLGRLYF | TRAV10 | TRAJ18 | TRAC | CASSESLSDTSTSTDTQYF | TRBV25-1 | TRBJ2-3 | TRBC2 | **iNKT** |

**Table 2.** MAIT cell TCR sequences further validated.

| Clone # | Donor ID | CDR3α | TRAV | TRAJ | TRAC | CDR3β | TRBV | TRBJ | TRBC |
|---|---|---|---|---|---|---|---|---|---|
| 1 | Donor 3 | CAVLDSNYQLIW | TRAV1-2 | TRAJ33 | TRAC | CSASGDREEIYEQYF | TRBV20-1 | TRBJ2-7 | TRBC2 |
| 2 | Donor 3 | CAVIDSNYQLIW | TRAV1-2 | TRAJ33 | TRAC | CSARDLGGRNFGETQYF | TRBV20-1 | TRBJ2-5 | TRBC2 |
| 3 | Donor 2 | CAVMDSNYQLIW | TRAV1-2 | TRAJ33 | TRAC | CASSEALGGGNQPQHF | TRBV6-1 | TRBJ1-5 | TRBC1 |
| 4 | Donor 2 | CAGMDSNYQLIW | TRAV1-2 | TRAJ33 | TRAC | CASSETSGSTDTQYF | TRBV6-4 | TRBJ2-3 | TRBC2 |
| 5 | Donor 2 | CAVRDSNYQLIW | TRAV1-2 | TRAJ33 | TRAC | CATSGGQPTDTQYF | TRBV15 | TRBJ2-3 | TRBC2 |

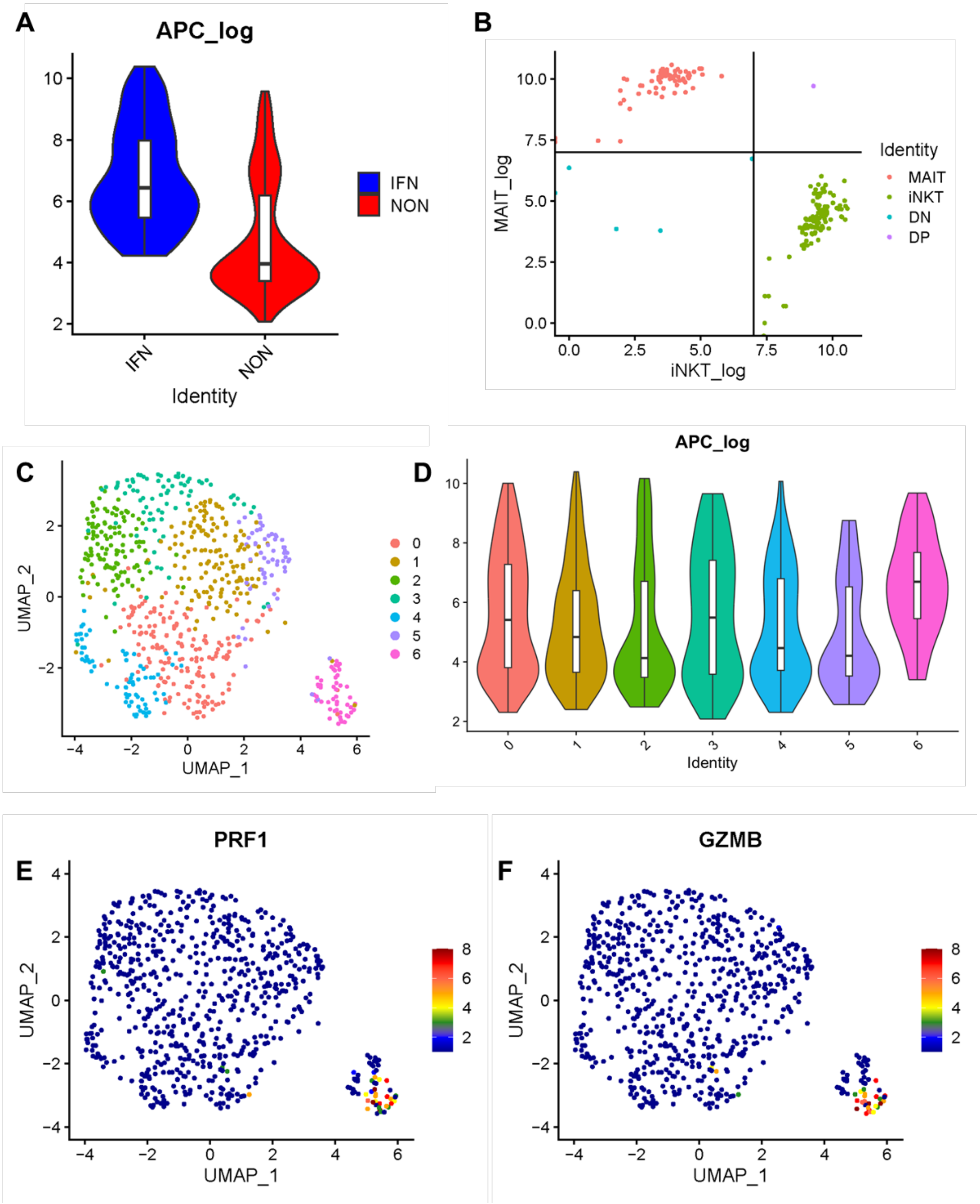


**Figure S5.** Single cell RNAseq results. (A) Anti-APC oligobarcode labeling counts is higher in IFNγ secretor population (blue) compared to non-secretor population (red). (B) Distributions of MR1- or CD1d-specific barcode reads for cells captured in either MR1 or CD1d nanovials. (C) Generated UMAP from single-cell RNA transcriptome data. (D) Anti-APC labeling signal, representing IFNγ levels, across the 6 identified clusters from the UMAP. (E) Perforin-1 (*PRF1*) gene expression overlaid on the generated UMAP. (F) Granzyme B (*GZMB*) gene expression overlaid on the generated UMAP.

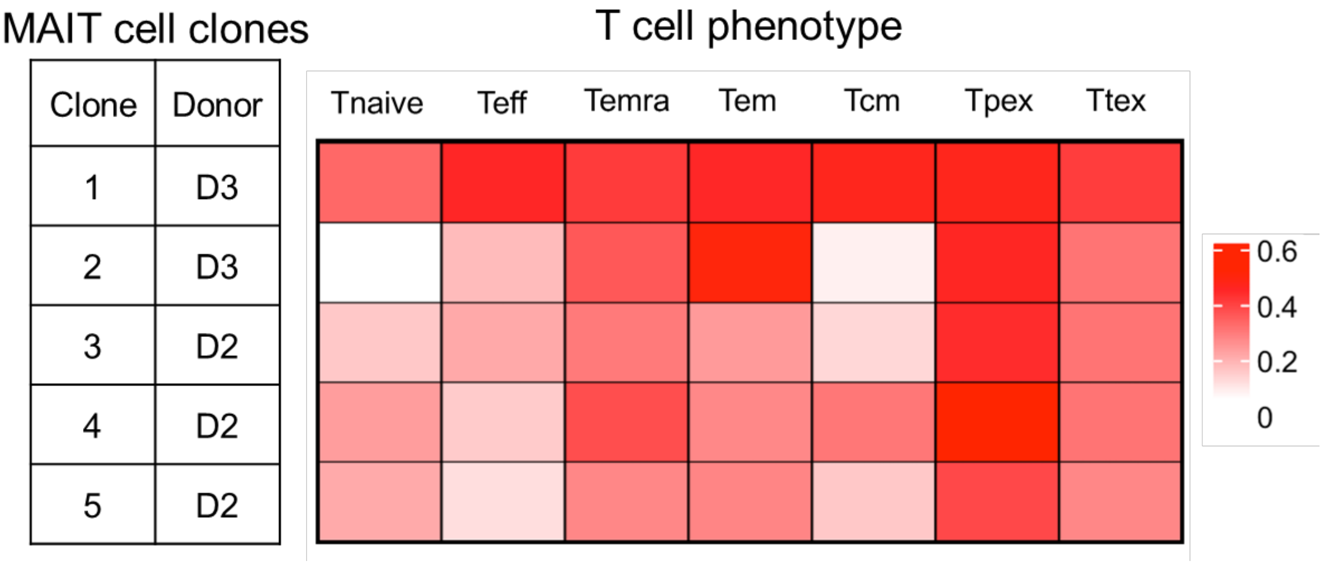


**Figure S6.** T cell phenotype based on gene expression for the five validated MAIT clones. Tnaive, naïve T cell; Teff, effector T cell; Temra, terminally differentiated effector memory T cell; Tem, effector memory T cell; Tcm, central memory T cell; Tpex, progenitor exhausted T cell; Ttex, terminally exhausted T cell. Scale bars indicate relative gene expression**.**

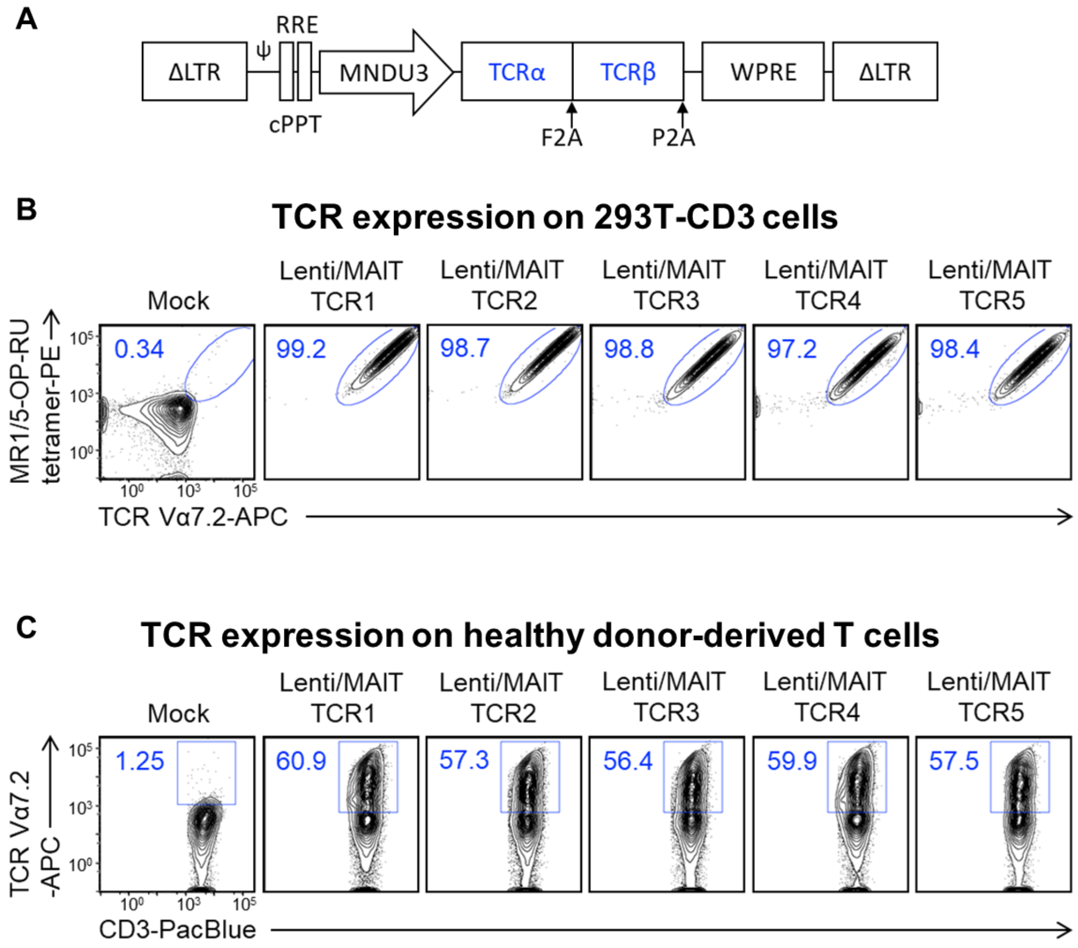


**Figure S7. Design and validation of the selected 5 MAIT TCRs.**

(A) Design of the lentivector. ΔLTR, self-inactivating long terminal repeats; MNDU3, internal promoter derived from the MND retroviral LTR U3 region; φ, packaging sequence; RRE, rev-responsive element; cPPT, central polypurine tract; WPRE, woodchuck hepatitis virus posttranscriptional regulatory element; F2A, foot-and-mouth disease virus 2 A. (B) FACS plots showing the successful expression of MAIT TCRs on 293T-CD3 cells. (C) FACS plots showing the successful expression of MAIT TCRs on healthy donor PBMC-derived T cells.

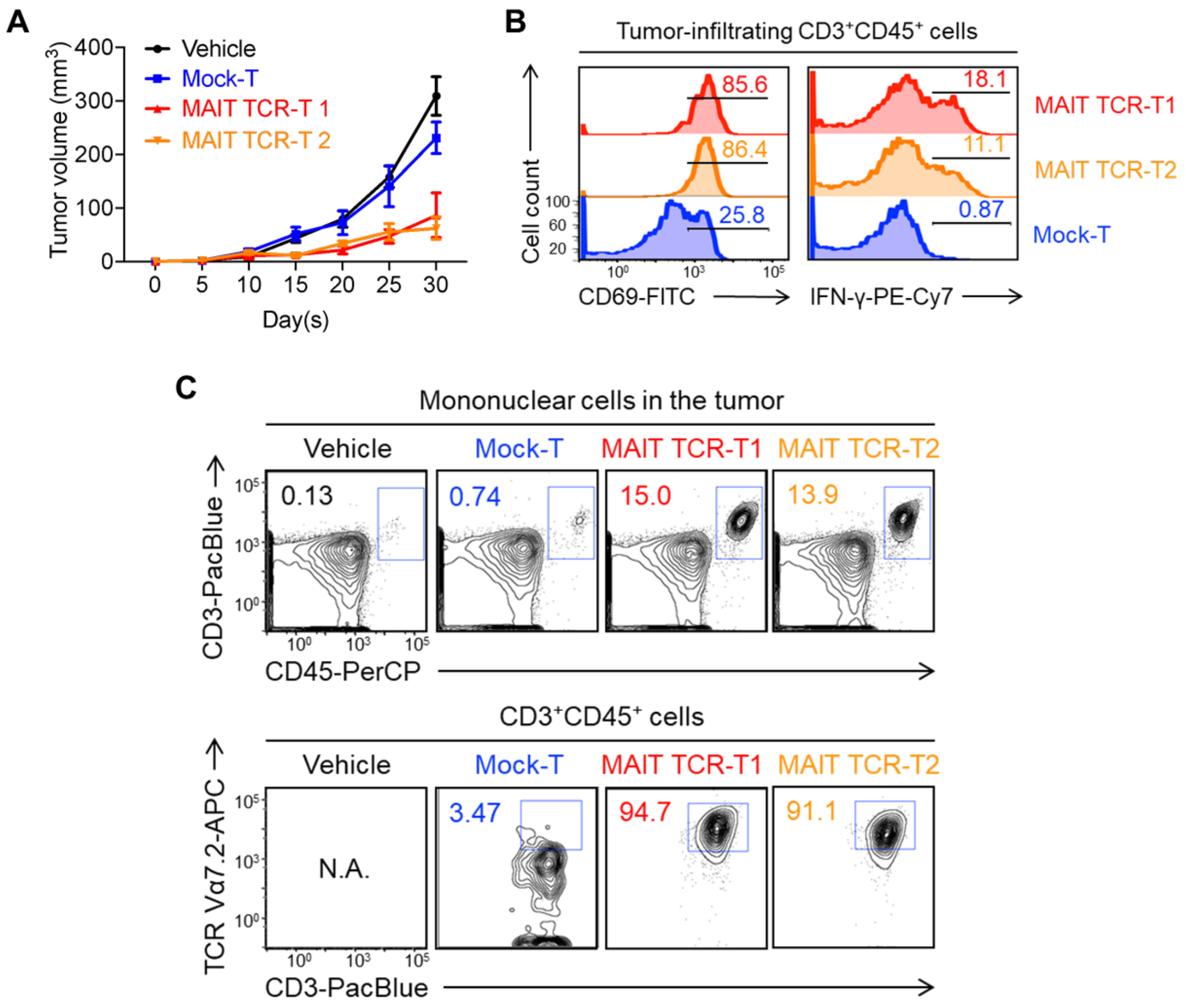


**Figure S8.** ***In vivo*** **validation using the top 2 MAIT TCRs using an A375 human melanoma xenograft mouse model.** (A) Tumor volume measurements over time (n = 5). (B) FACS plots showing the CD69 and IFNγ intracellular staining of the indicated tumor-infiltrating T cells. (C) FACS plots showing the detection of tumor-infiltrating T cells and the MAIT TCR-engineered T cells.
